# Harmful DNA:RNA hybrids are formed *in cis* and in a Rad51-independent manner

**DOI:** 10.1101/2020.04.20.047134

**Authors:** Juan Lafuente-Barquero, Marta San Martin-Alonso, Belén Gómez-González, Andrés Aguilera

## Abstract

DNA:RNA hybrids constitute a well-known source of recombinogenic DNA damage. The current literature is in agreement with DNA:RNA hybrids being produced co-transcriptionally by the invasion of the nascent RNA molecule produced *in cis* with its DNA template. However, it has also been suggested that recombinogenic DNA:RNA hybrids could be facilitated by the invasion of RNA molecules produced *in trans* in a Rad51-mediated reaction. Here, we tested the possibility that such DNA:RNA hybrids produced *in trans* constitute a source of recombinogenic DNA damage taking advantage of Rad51-independent single-strand annealing recombination assays and new constructs designed to induce expression of mRNA transcripts *in trans* in the yeast *Saccharomyces cerevisiae.* We show that unscheduled and recombinogenic DNA:RNA hybrids are formed in *cis* during transcription and in a Rad51-independent manner.

## INTRODUCTION

R loops are structures formed by a DNA:RNA hybrid and the complementary displaced single stranded DNA (ssDNA). They were observed naturally as programmed events in specific genomic sites such as the S regions of Immunoglobulin genes in mammals or mitochondrial DNA (Chang, Hauswirth, & Clayton, 1985; Garcia-Muse & Aguilera, 2019; Yu, Chedin, Hsieh, Wilson, & Lieber, 2003), where they play specific functions by promoting class switch recombination or DNA replication, respectively; but also as unscheduled nonprogrammed structures upon dysfunction of RNA binding proteins involved in the assembly or processing and export of the protein-mRNA particle (mRNP) such as the THO complex or the SRSF1 splicing factor (Huertas & Aguilera, 2003; X. Li & Manley, 2005). Also, they have been inferred in the rDNA regions of the bacterial chromosome upon Topo1 inactivation (Drolet et al., 1995). Accumulated evidence indicates that R loops are detected from yeast to humans in many transcribed regions of the eukaryotic genome in wild-type cells as well as in cells defective in several metabolic processes covering from RNA processing to DNA replication and repair or cells deficient in specific chromatin factors (Bhatia et al., 2014; Garcia-Muse & Aguilera, 2019; Garcia-Rubio et al., 2015; Herrera-Moyano, Mergui, Garcia-Rubio, Barroso, & Aguilera, 2014; Mischo et al., 2011; Paulsen et al., 2009; Schwab et al., 2015). The biological consequences of such R loop structures are diverse and include replication stress, DNA breaks and genome instability that can be detected as hyperrecombination, plasmid loss or gross chromosomal rearrangements (Garcia-Muse & Aguilera, 2019). Indeed, DNA:RNA hybrids have been inferred by their potential to induce DNA damage and recombination, but they can also be directly detected via different methodologies. These include electrophoresis detection after nuclease treatment, bisulfite mutagenesis or either in situ immunofluorescence or DNA:RNA Immuno-Precipitation (DRIP) using the S9.6 anti-DNA:RNA monoclonal antibody (Garcia-Muse & Aguilera, 2019).

The increasing number of reports showing R loop accumulation in different organisms from bacteria to human cells, and the relevance of their functional consequences, whether on genome integrity, chromatin structure and gene expression suggest that most DNA:RNA hybrids are compatible with a co-transcriptional formation (Garcia-Muse & Aguilera, 2019). This is consistent with the idea that it is the RNA produced *in cis* who invades the duplex DNA, a reaction that can be facilitated by increasing the negative supercoiling of DNA as well as by nicking the DNA template (Roy, Zhang, Lu, Hsieh, & Lieber, 2010). The evidence of DNA:RNA hybrid formation at breaks has matured in the last years (Cohen et al., 2018; D’Alessandro et al., 2018; L. Li et al., 2016; Ohle et al., 2016; Teng et al., 2018; Yasuhara et al., 2018) although the source and role of such hybrids remains still controversial (Aguilera & Gomez-Gonzalez, 2017; Puget, Miller, & Legube, 2019). Of note, genome-wide mapping results have been interpreted in diverse manners by different labs. Whereas some claim that DNA:RNA hybrids detected around DNA breaks mostly accumulate at transcribing sites (Cohen et al., 2018), in agreement with their co-transcriptional formation, others suggest that there is no preference for DNA:RNA hybrids to form at transcribed loci (D’Alessandro et al., 2018), implying an scenario in which DNA:RNA hybrids at breaks sites would form either *de novo* or with RNAs produced *in trans*.

DNA:RNA hybrids may be formed *in trans* as intermediates in the course of ribonucleoprotein-mediated reactions such as telomerase and CRISPR-Cas9 ribonucleoprotein involved in specific reactions (Collins, 2000; Jinek et al., 2012). DNA:RNA hybrids can also be formed *in vitro* with the aid of the bacterial strand exchange protein RecA (Kasahara, Clikeman, Bates, & Kogoma, 2000; Zaitsev & Kowalczykowski, 2000). They have also been reported to have regulatory roles in gene expression when formed by long non-coding RNAs (lncRNAs) at *in trans* loci such as the cases of the GAL lncRNA in yeast (Cloutier et al., 2016) or the APOLO lncRNA in plants (Ariel et al., 2020). In summary, despite the accumulating evidence that *in vivo* DNA:RNA hybrids formed *in cis* constitute a threat for genome stability, an open question is whether DNA:RNA hybrids also form *in trans* as a potential source of recombinogenic DNA damage. To our knowledge, this has only been addressed in the yeast *Saccharomyces cerevisiae* (Wahba, Gore, & Koshland, 2013). By S9.6 immunofluorescence (IF) and a yeast artificial chromosome-based genetic assay that measures gross chromosomal rearrangements, it was inferred that DNA:RNA hybrids could be formed *in trans* by a reaction catalyzed by the eukaryotic strand-exchange protein Rad51 (Wahba et al., 2013).

Nevertheless, the fact that the detected gross chromosomal rearrangements could depend on Rad51 and that the S9.6 antibody can also recognize dsRNAs (Hartono et al., 2018; Konig, Schubert, & Langst, 2017; Silva, Camino, & Aguilera, 2018), prompted us to address this question using a different approach. Using Rad51-independent recombination assays in which the initiation region could be unambiguously delimited, we provide genetic evidence that DNA:RNA hybrids compromising genome integrity are formed in *cis* and in a Rad51-independent manner.

## RESULTS

### A new genetic assay to detect recombinogenic DNA:RNA hybrids *in trans*

We developed a new genetic assay to infer the formation of recombinogenic DNA:RNA hybrids *in trans.* It is based on two plasmids, one containing the recombination system and the *LacZ* gene *in cis (GL-LacZ* recombination system), and another one providing the *in trans LacZ* transcripts (*tet_p_:LacZ*) (Figure 1). The bacterial *LacZ* gene consists of a 3-Kb sequence with high G+C content previously reported to be hyper-recombinant and difficult to transcribe in DNA:RNA hybrid-accumulating strains, such as *tho* mutants (Chavez, Garcia-Rubio, Prado, & Aguilera, 2001).

**Figure 1.**
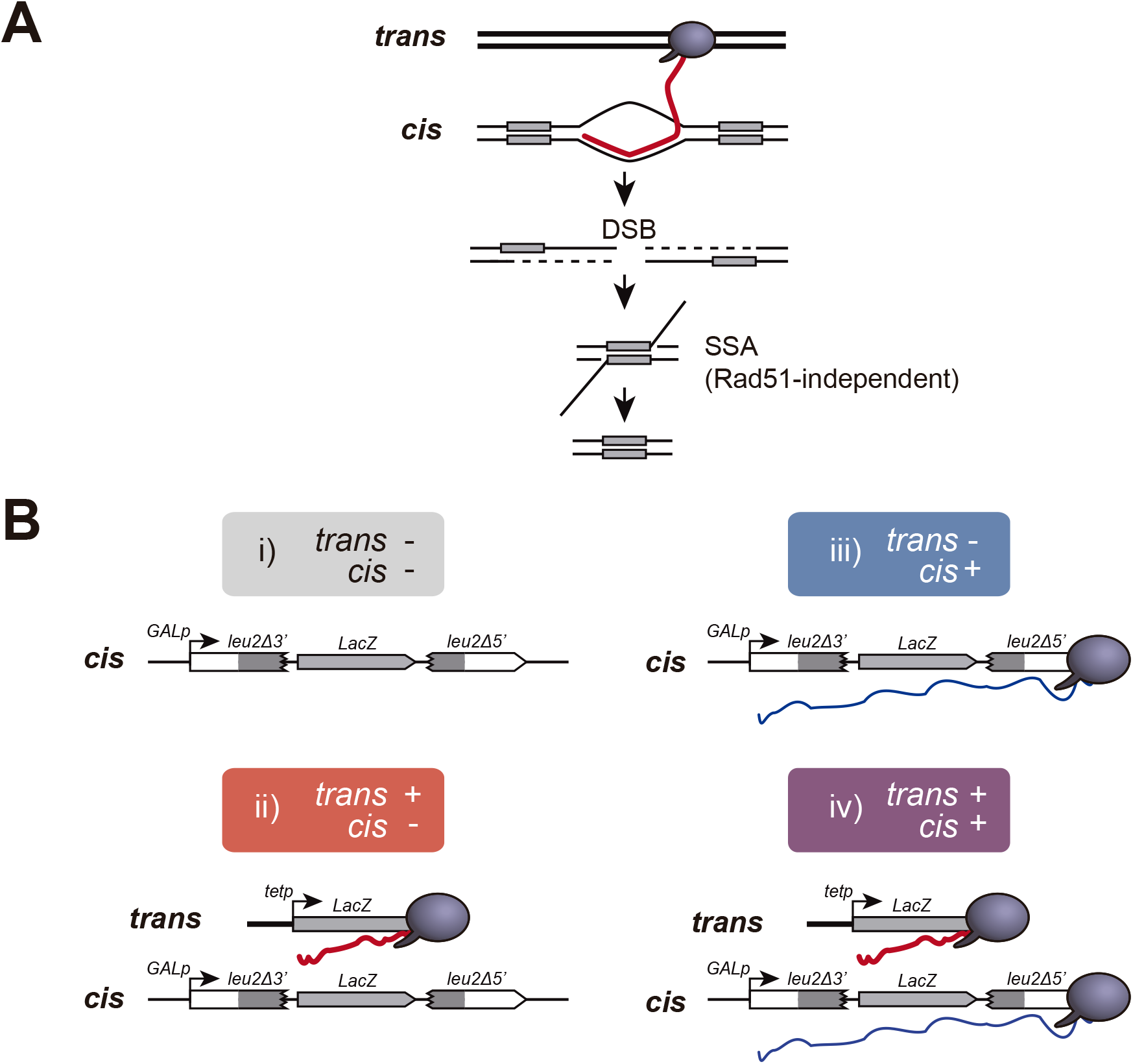
A new genetic assay to detect recombinogenic DNA:RNA hybrids *in trans.* **(A)** DSBs induced in between direct repeats by DNA:RNA hybrids putatively formed with RNA produced *in trans* would be repaired by Rad51-independent Single-Strand Annealing (SSA) causing the deletion of one of the repeats. **(B)** Schematic representation of the recombination assay to study the recombinogenic potential RNA produced by transcription (Trx) *in cis* or *in trans*. Four combinations were studied: i) no transcription, with GL-*LacZ* construct turned transcriptionally off (2% glucose) and an empty plasmid; ii) transcription *in trans*, with GL-*LacZ* construct turned transcriptionally off (2% glucose) and the *tetp:LacZ* construct; iii) transcription *in cis*, with GL-*LacZ* construct turned transcriptionally on (2% galactose) and an empty plasmid; and iv) transcription *in cis* and *in trans*, with GL-*LacZ* construct turned transcriptionally on (2% galactose) and the *tet_p_:LacZ* construct.

The GL-*LacZ* recombination system is a *leu2* direct-repeat construct carrying the *LacZ* gene in between and under the *GAL1* inducible promoter so that this construct is transcribed as a single RNA unit driven from the *GAL1* promoter (Piruat & Aguilera, 1998). Single-Strand Annealing (SSA) events cause the deletion of the *LacZ* sequence and one of the *leu2* repeats leading to Leu+ recombinants in a Rad51-independent manner (Figure 1A). To provide *LacZ* transcripts *in trans*, we did a fusion construct containing the complete bacterial *LacZ* gene sequence under the doxycycline-inducible *tet* promoter (*tetp:LacZ*). As a control of no expression *in trans*, we used transformants with an empty plasmid to avoid any possible effect from leaky transcription from the *tet* promoter in the presence of doxycycline.

Yeast strains carrying both GL-*LacZ* recombination system and the *tet_p_:LacZ* construct were used to assay SSA annealing events in the four different possible conditions: i) no transcription, with GL-*LacZ* construct turned transcriptionally off (2% glucose) and an empty plasmid; ii) transcription *in trans*, with GL-*LacZ* construct turned transcriptionally off (2% glucose) and the *tetp:LacZ* construct; iii) transcription *in cis*, with GL-*LacZ* construct turned transcriptionally on (2% galactose) and an empty plasmid; and iv) transcription *in cis* and *in trans*, with GL-*LacZ* construct turned transcriptionally on (2% galactose) and the *tetp:LacZ* construct (Figure 1B).

### RNA is not a spontaneous source of recombinogenic DNA damage *in trans*

The analysis of recombination in wild-type cells revealed that whereas transcription *in cis* elevated the frequency of recombination threefold, transcription *in trans* driven from the *tetp:LacZ* construct had no effect on recombination (Figure 2A). These results already suggest that homologous transcripts coming from a different locus do not represent a detectable source of genetic instability in wild-type conditions and thus argue against the hypothesis that spontaneous DNA:RNA hybrids could be formed with mRNAs transcribed *in trans.* However, it is known that mRNA coating protects DNA from co-transcriptional RNA hybridization. Thus, we wondered if *in trans* transcripts could induce recombination in mRNP-defective mutants such as those of the THO complex. Hence, we performed our experiments in *mft1*Δ and *hpr1*Δ mutant strains. *mft1*Δ and *hpr1*Δ enhanced recombination slightly when transcription *in cis* was switched off, likely as a consequence of leaky transcription form the *GAL1* promoter in glucose. More significantly and in agreement with previous reports (Chavez et al., 2000), recombination frequencies rocketed when transcription was stimulated *in cis*. However, transcription activation *in trans* did not enhance recombination, as it would be expected if additional DNA:RNA hybrids could form with RNA produced *in trans.*

**Figure 2.**
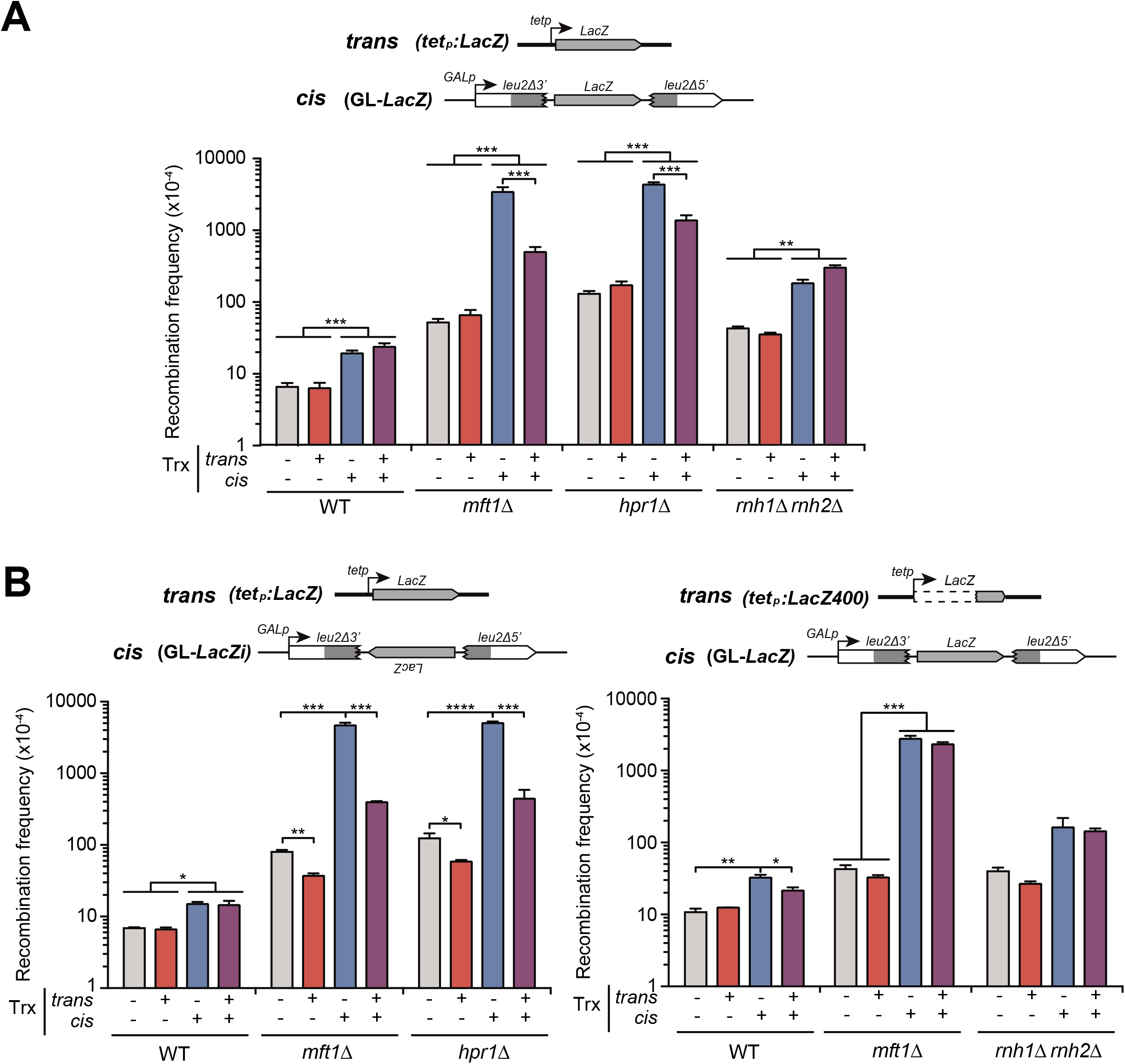
Analysis of the effect on genetic recombination of RNA produced *in cis* or *in trans.* **(A)** Recombination analysis in WT (W303), *rnh1Δ rnh201*Δ (HRN2.10C), *mft1*Δ (WMK.1A) and *hpr1*Δ (U678.4C) strains carrying *GL-LacZ* plasmid system (pRS314-GL-*LacZ*) plus either the empty vector (pCM190) or the same vector carrying the *LacZ* gene (pCM179). **(B)** Recombination analysis in WT (W303), *mft1*Δ (WMK.1A) and *hpr1*Δ (U678.4C) strains carrying plus either the empty vector pCM190 or the same vector carrying the sequence of the *LacZ* gene (pCM179). **(C)** Recombination analysis in WT (W303), *rnh1Δ rnh201*Δ (HRN2.10C) and *mft1*Δ (WMK.1A) strains carrying *GL-LacZ* plasmid system (pRS314-GL-*LacZ*) plus either the empty vector pCM190 or the same vector carrying the last 400 bp from the 3’ end of the *LacZ* gene (pCM190:*LacZ400*). In all panels, average and SEM of at least three independent experiments consisting in the median value of six independent colonies each are shown. *, p≤0.05; **, p≤0.01; ***, p≤0.001; ****, p≤0.0001. (Student’s t-test).

Instead, under conditions of high transcription *in cis,* transcription *in trans* led to a partial suppression of the hyper-recombination observed by only *in cis* transcription. The reason for such a suppression might involve the potential ability of the mRNA produced *in trans* to interfere with transcription occurring *in cis* at the *GL-LacZ* construct. Given that a DNA:RNA hybrid produced in the template DNA strand can impair transcription elongation (Tous & Aguilera, 2007), one possibility was that this interference is mediated by DNA:RNA hybrids formed *in trans* with the transcribed strand of the GL-*LacZ* construct. However, this possibility was ruled out by the fact that transcription *in trans* also led to a reduction of the hyper-recombination when we used an alternative recombination system (GL*-LacZi*), in which the *LacZ* sequence was inverted so that the *LacZ* transcript produced *in trans* would not be able to anneal with the transcribed strand of the GL-*LacZi* system (Figure 2B). Furthermore, in this case the suppression was stronger and was also observed in glucose, when transcription *in cis* was off. This could be explained because, in this scenario, the RNA produced *in trans* is complementary to the mRNA produced *in cis*. Consequently, they can hybridize together forming a dsRNA that would preclude the possibility to form DNA:RNA hybrids at the *GL-LacZi* construct.

RNAses H efficiently degrade the RNA moiety of DNA:RNA hybrids. Thus, to favor the potential DNA:RNA hybrid accumulation *in trans* we used cells lacking also both RNAses H1 and H2 and we determined the impact on recombination. Figure 2A shows that *rnh1Δ rnh201*Δ cells elevated the recombination frequency when transcription was stimulated *in cis.* Importantly, the recombination frequencies were not altered by producing transcripts *in trans*, arguing again against the recombinogenic potential of putative DNA:RNA hybrids formed *in trans*.

Since transcription from the long *LacZ* gene is inefficient and leads to unstable RNA products, particularly in *tho* mutants (Chavez et al., 2001), we made a new construct with only the last 400 bp of *LacZ (tet_p_:LacZ400)* (Figure 2C). In this case, recombination frequencies were not significantly affected by *in trans* transcription in any of the strains or conditions tested further arguing against mRNA produced *in trans* as a possible source of recombinogenic DNA:RNA hybrids.

Finally, to confirm that the same results were obtained in chromosome *loci* and not only in plasmid-born DNA sequences, we integrated the GL-*LacZ* system in the chromosome. As shown in Figure 3, we observed again that mRNA production *in trans* had no effect on recombination, neither in wild-type cells nor in the *tho* mutant *hpr1Δ.* Furthermore, the results were similar after RNase H overexpression. Hence, altogether, these results argue that, in contrast to mRNA produced *in cis*, RNA produced at a particular *locus* does not lead to recombinogenic DNA damage at regions located *in trans*.

**Figure 3.**
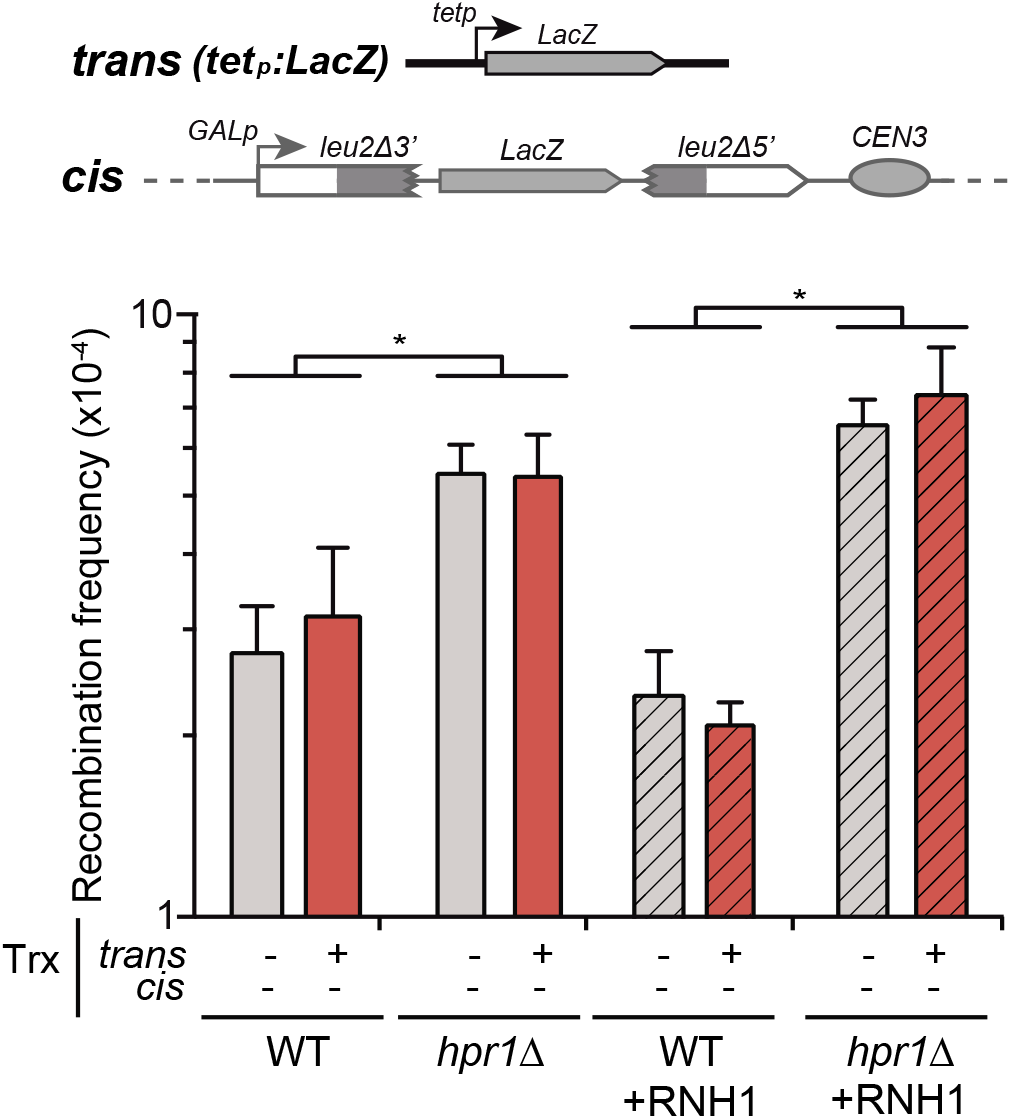
Analysis of the effect on genetic recombination of RNA produced *in trans* of chromosome III with or without RNAse H overexpression. Recombination analysis in WT (WGLZN), *hpr1*Δ (HGLZN) strains carrying the *GL-LacZ* recombination system integrated in chromosome III. Strains were transformed with empty vector pCM190 or the same vector carrying the *LacZ* gene (pCM179) plus either plasmid pCM184 (empty vector) or the same vector carrying the RNH1 gene (RNH1). Average and SEM of at least three independent experiments are shown consisting in the median value of six independent colonies each. *, p≤0.05. (Student’s t-test).

### Rad51 is not required for DNA:RNA hybridization

We next wondered about the possible role of the recombination protein Rad51 in DNA:RNA hybridization. To examine this, in particular in relation to hybrids potentially formed *in trans*, we analyzed recombination frequencies in our direct-repeat assays in *hpr1*Δ cells in which *in cis* transcription was switched off, under conditions in which an homologous RNA with the potential to hybridize with the intervening sequence of repeat construct was or was not generated *in trans* (Figure 4A). It is important to remark that the recombination events detected in our assays are deletions occurring by SSA between direct repeats, which do not require Rad51 (Pardo, Gomez-Gonzalez, & Aguilera, 2009). Indeed, in agreement with SSA annealing being Rad51-independent, *RAD51* deletion caused no significant changes in the recombination frequencies in our assay. Thus any conclusion about Rad51-dependency or independency of the hybridization inferred from our assay is not contaminated by a possible direct role of Rad51 in the event we are studying. Importantly, we observed no differences when *RAD51* was deleted in *hpr1*Δ cells even when the *LacZ* sequence was expressed *in trans*. This result argues against Rad51 facilitating or impeding the formation of DNA:RNA hybrids *in trans.*

**Figure 4.**
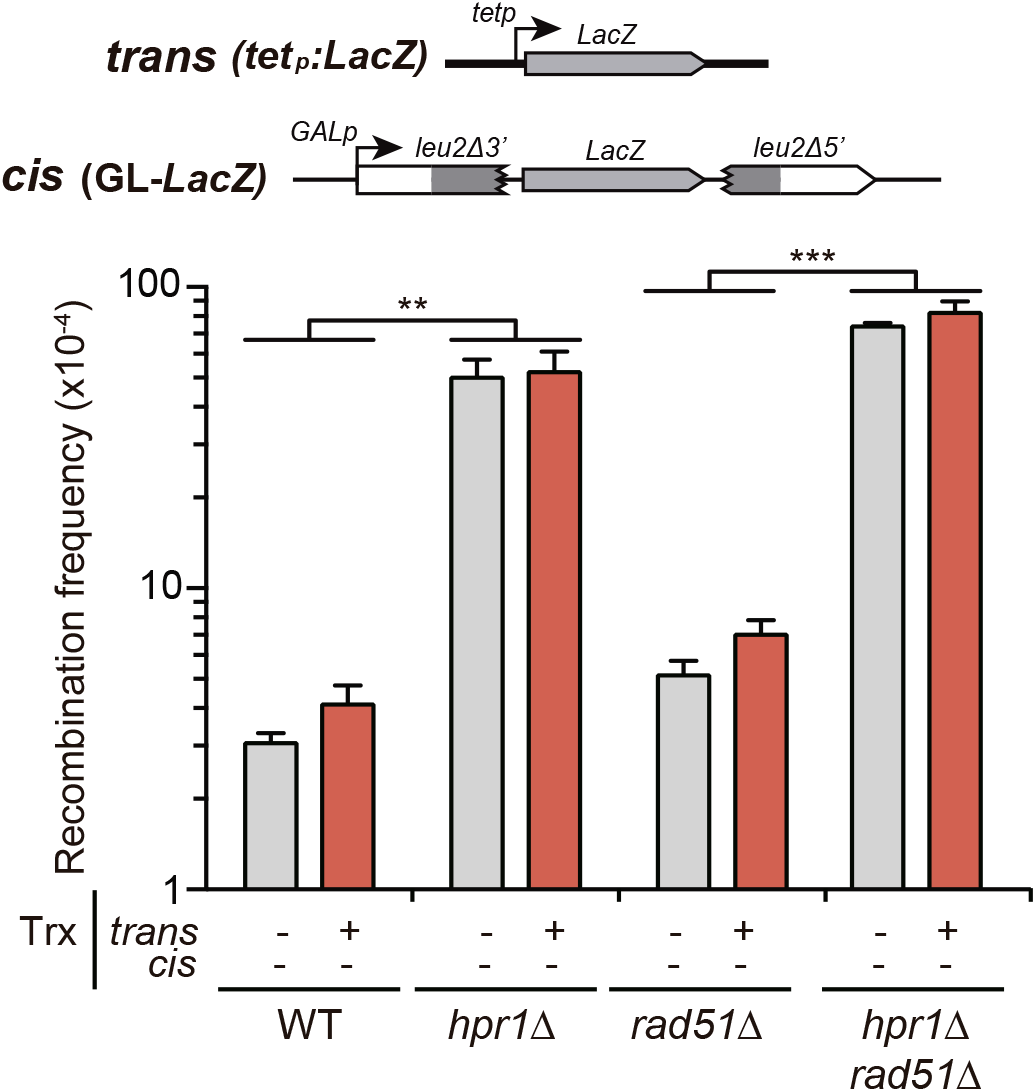
Analysis of the effect on genetic recombination of RNA produced *in trans* with or without Rad51. Recombination analysis in WT (W303), *hpr1*Δ (U678.1C), *rad51*Δ (WSR51.4A) and *hpr1Δ rad51*Δ (HPR51.15A) strains carrying *GL-LacZ* plasmid system (pRS314-GL-*LacZ*) plus either the empty vector pCM190 or the same vector carrying the *LacZ* gene (pCM179). Average and SEM of at least three independent experiments consisting in the median value of six independent colonies each are shown. **, p≤0.01; ****, p≤0.0001. (Student’s t-test). (

We then wondered whether the formation of known recombinogenic DNA:RNA hybrids formed *in cis,* such as those reported in the *hpr1*Δ mutant, requires Rad51. For this purpose, we deleted *RAD51* and studied the effect in the strong hyper-recombination phenotype of *hpr1Δ,* which has been observed in multiple direct-repeat systems in which recombinants also arise by Rad51-independent SSA, such as the LYΔNS system consisting in two truncated *leu2* direct-repeats under the *LEU2* promoter with a 3.7-Kb sequence in between (Prado, Piruat, & Aguilera, 1997). As shown in Figure S1, *hpr1*Δ led to a 33-fold increase in recombination in the absence of Rad51, similarly to what was previously reported in the presence of Rad51 (Gomez-Gonzalez, Felipe-Abrio, & Aguilera, 2009; Prado et al., 1997). This result clearly indicates that the *in cis* DNA-RNA hybrid-mediated hyper-recombination phenotype is actually independent on Rad51.

In parallel, we studied the formation of Rad52 foci, a marker of recombinogenic DNA breaks (Lisby, Rothstein, & Mortensen, 2001) as well as the effect of AID overexpression to enhance the recombinogenic potential of R loops (Gomez-Gonzalez & Aguilera, 2007) and RNase H overexpression to remove DNA:RNA hybrids (Figure 5A). In agreement with the role of the THO complex in R loop prevention, *hpr1*Δ caused an increase in Rad52 foci that was enhanced by AID overexpression and suppressed by RNase H overexpression, as previously reported (Alvaro, Lisby, & Rothstein, 2007; Garcia-Pichardo et al., 2017; Wellinger, Prado, & Aguilera, 2006). By contrast, the accumulation of Rad52 foci observed in *rad51*Δ cells was not affected by either AID or RNase H overexpression. This result argues that R loops are not the cause for the genetic instability observed in the absence of Rad51. Given the role of Rad51 in recombination upstream of Rad52, the accumulation of Rad52 foci in *rad51*Δ cells is rather likely due to the accumulation of unrepaired recombination intermediates, as previously suggested (Alvaro et al., 2007). Importantly, *hpr1Δ rad51*Δ cells showed a similar result, further supporting that the accumulation of recombinogenic damage in *hpr1*Δ is independent on Rad51. We next directly measured DNA:RNA hybrid accumulation by immunodetection with the S9.6 antibody on metaphase spreads. Figure 5B illustrates that the number of cells with S9.6 positive signal was similar in *hpr1*Δ and in *hpr1Δ rad51*Δ cells. Altogether, these results demonstrate that the Rad51 protein is not required for the DNA:RNA hybrid formation previously reported in THO mutants.

**Figure 5.**
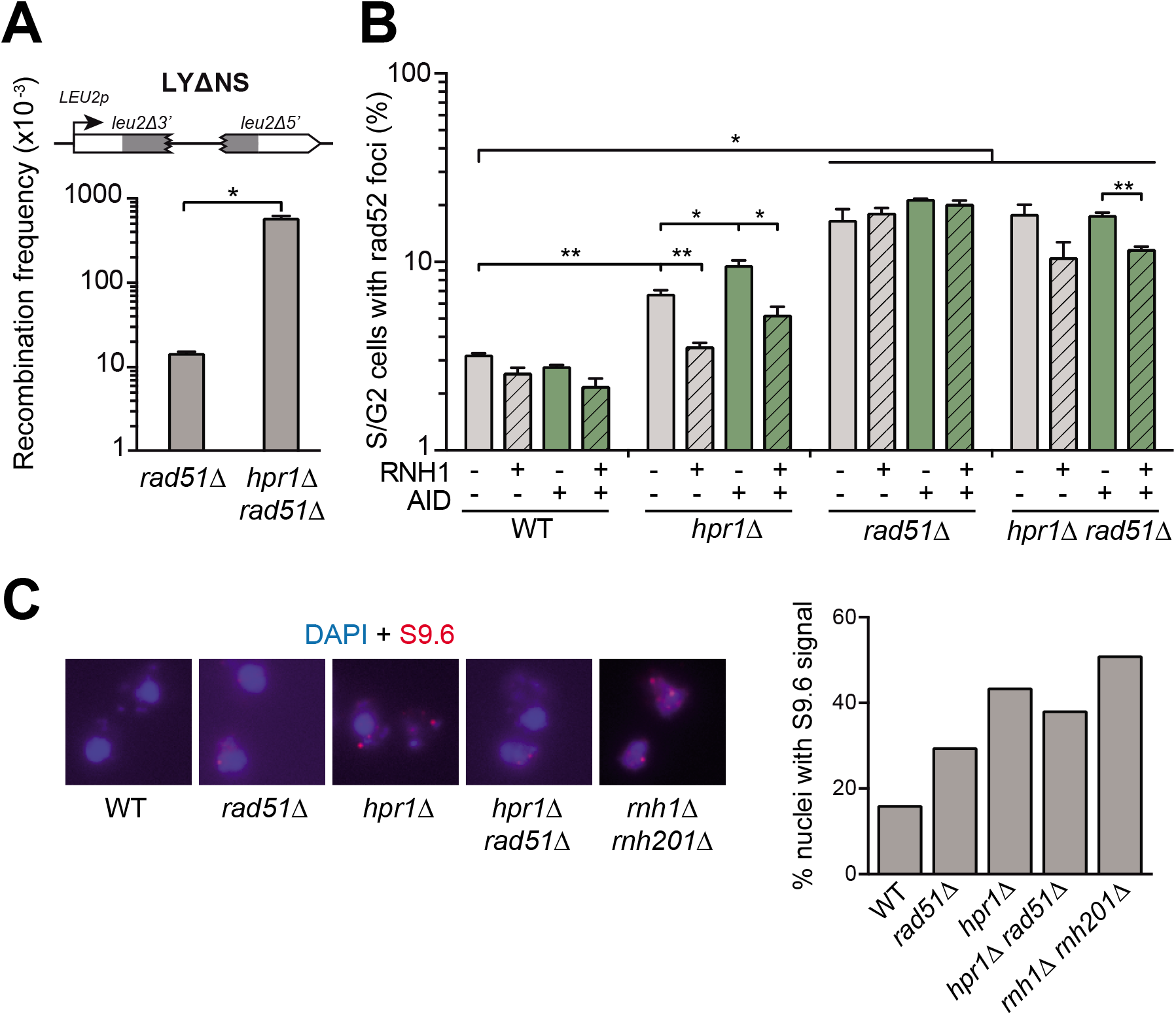
The increased genetic instability and DNA:RNA hybrids of *hpr1*Δ is independent on Rad51. **(A)** Recombination analysis in *rad51*Δ (AYBY-7B) and *hpr1Δ rad51*Δ (AYBY-9B) strains carrying LYΔNS plasmid system (pRS316-LYΔNS). Average and SEM of at least three independent experiments consisting in the median value of six independent colonies each are shown. *, p≤0.05; **, p≤0.01; ***, p≤0.001. (Student’s t-test). **(B)** Spontaneous Rad52-YFP foci formation in WT (W303), *hpr1*Δ (U678.1C), *rad51*Δ (WSR51.4A) and *hpr1Δ rad51*Δ (HPR51.15A) strains carrying the empty vectors pCM184 and pCM189, or a combination of both carrying the RNH1 or AID genes as indicated in the legend. Average and SEM of at least three independent experiments are shown. *, p≤0.05; **, p≤0.01. (Student’s t-test). **(C)** Representative images and median value of the percent of the total nuclei scored that stained positively for DNA:RNA hybrids in chromatin spreads stained with the S9.6 antibody in WT (W303), *hpr1*Δ (U678.1C), *rad51*Δ (WSR51.4A), *hpr1*Δ *rad51*Δ (HPR51.15A) and RNH-R (*rnh1*Δ *rnh201*Δ) strains.

## DISCUSSION

We have devised a new genetic assay to infer whether the source of DNA:RNA hybrids compromising genome integrity could potentially come from RNAs produced *in trans.* To reach this conclusion, we used an SSA assay. It is well established that SSA events are Rad51-independent; they do not require strand-exchange, but just annealing between resected single-stranded DNA (ssDNA) for which the action of Rad52 is sufficient (Figure 1A) (Pardo et al., 2009). Our constructs show that, in contrast to the RNA produced *in cis* at the site where SSA occurs, an RNA produced *in trans* does not induce an increase in recombination. Importantly, recombination is not induced by *in trans* RNA production even when the major DNA:RNA removal machinery is absent *(rnh1Δ rnh201*Δ mutant) or when the RNA coating functions preventing DNA:RNA hybrid formation are impaired (*tho* mutants), arguing against the idea that harmful DNA:RNA hybrids could spontaneously form *in trans.* Putative DNA:RNA hybrids formed *in trans* would be expected to further increase recombination levels. Instead, the simultaneous induction of transcription *in cis* and *in trans* (Figure 2A) reduced the strong hyper-recombinogenic effect of *tho* mutants. The fact that this suppressor effect was augmented when one of the *LacZ* sequences was inverted (Figure 2B) and prevented by a shorter *LacZ* construct (Figure 2C), which was reported to be more stable in *tho* mutant backgrounds (Chavez et al., 2001), suggests that the free RNA itself, and not in the form of DNA:RNA hybrids formed at the template DNA strand, play some role in preventing the hyper-recombination, likely because stable RNAs can interfere with transcription at an homologous locus.

DNA:RNA hybrids formed by an RNA produced *in trans* were previously suggested to threaten genome integrity based on the results obtained with a yeast artificial chromosome and an homologous region placed at chromosome III, which transcription was inducible (Wahba et al., 2013). Recombination involving multiple substrates was first reported in *S. cerevisiae*, in which an induced-DSB triggered recombination between two other homologous fragments at different chromosomes (Ray, Machin, & Stahl, 1989). Tri-parental recombination assays have been successfully used since then to define specific features of the HR reaction as well as for studies of Break-Induced Recombination (BIR) or translocations and chromosomal rearrangements occurring between ectopic regions (Pardo & Aguilera, 2012; Piazza, Wright, & Heyer, 2017; Ruiz, Gomez-Gonzalez, & Aguilera, 2009). However, such events are not the most adequate to infer recombination initiation unless this has been artificially designed (as is the case of an HO-induced DSB). Hence, the assay used to infer the potential of *trans* DNA:RNA hybrids to induce genetic instability (Wahba et al., 2013) relied on an RNA fragment produced at a (first) DNA region that could form a DNA:RNA hybrid with a (second) ectopic homologous DNA region that would promote its deletion or loss, leading to a genetically detectable phenotype. Thus, this assay does not exclude the possibility that the RNA forms the hybrid *in cis* inducing subsequently a DSB that would stimulate the recombination events studied (Figure 6). Indeed, this event would demand the action of Rad51 for DNA strand invasion, consistent with the results obtained (Wahba et al., 2013). Therefore, rather than implying that Rad51 is required for the RNA to invade *in trans* the second DNA sequence, the increased genetic instability obtained suggests that the 3’ end of the DNA break induced by the DNA:RNA hybrid formed at the first site who makes the invasion (Figure 6).

**Figure 6.**
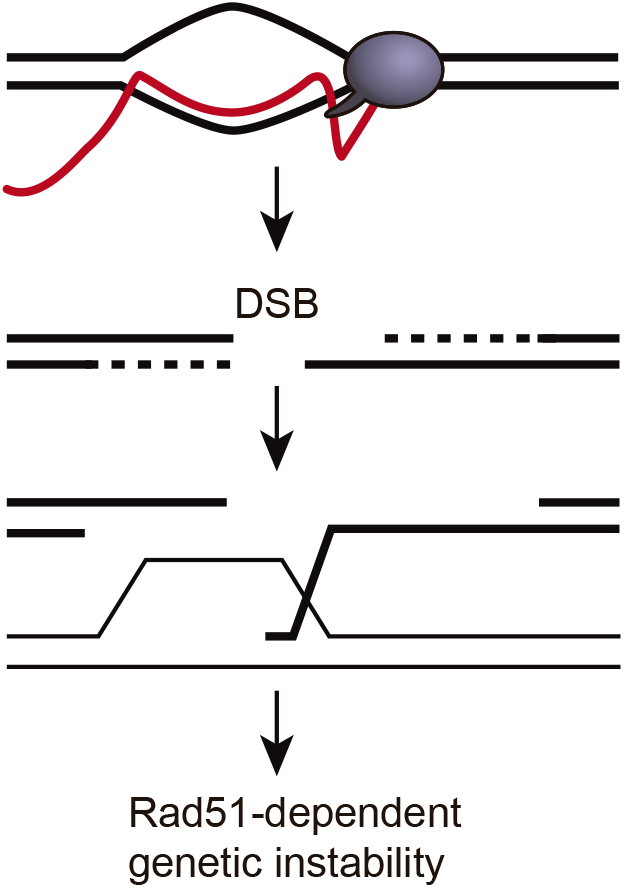
A model to explain how DNA:RNA hybrids could induce Rad51-dependent genetic instability *in trans.* DNA:RNA hybrid produced *in cis* can induce a DSB in the same sequence. The 3’ end of such a DSB could invade an ectopic homologous sequence and destabilize it. This DNA strand invasion event would require Rad51. In this model, genetic instability caused by hybrids *in trans* would be Rad51-dependent without the need of invoking a Rad51-mediated DNA-RNA hybrid formation *in trans*.

Our assays involve two *leu2* homologous repeats that recombine by Rad51-independent SSA. Indeed, as expected, *RAD51* deletion caused no decrease in the observed recombination frequencies in our assay (Figure 4). Recombination between the *leu2* repeats could be originated by either a DNA:RNA hybrid *in cis* or by a DSB occurring in between the repeats, or as suggested previously for *tho* mutants, by a bypass mechanism involving template switching (Gomez-Gonzalez et al., 2009). In our case, however, we show that the hyper-recombinogenic potential of DNA:RNA hybrids is Rad51-independent (Figure 4).

Similarly, a DSB occurring at the locus where the RNA *in trans* was generated could give rise to Leu+ recombinants in our assay. However, such recombination events would be Rad51-dependent, as they will require a Rad51-dependent invasion into the GL-*LacZ* construct (Figure 6). Hence, the Leu+ recombinants obtained in *rad51*Δ mutant cells (Figure 4) can only be explained by Rad51-independent events occurring *in cis*, at the GL-*LacZ* construct. Strikingly, the fact that we detected no significant increase in Leu+ recombinants by inducing transcription *in trans,* either in *RAD51* or *rad51*Δ backgrounds rules out the possibility that recombinogenic DNA:RNA hybrids form *in trans* in our assay. It was previously shown that S9.6 signal detected by IF was reduced by *rad51*Δ in metaphase spreads (Wahba et al., 2013). By contrast, we detected S9.6 signal in metaphase spreads of the *hpr1*Δ mutant of the THO complex in both *RAD51* and *rad51*Δ backgrounds (Figure 5). The uncertainty about the identity of the structures detected by IF using the S9.6 antibody, which also recognizes dsRNA (Hartono et al., 2018; Konig et al., 2017; Silva et al., 2018), and the possibility that chromosomal spreads could preferentially visualize the rDNA regions, in which high levels of dsRNA structures formed by the rRNAs, makes difficult to make conclusions on S9.6 IFs in this case.

Thus, we have found no evidence for a Rad51-facilitated strand invasion from RNAs produced *in trans*. Further arguing against any major role of this recombinase in R loop metabolism or function, none of the so far reported DNA:RNA hybrid interactomes has identified RAD51 (Cristini, Groh, Kristiansen, & Gromak, 2018; Nadel et al., 2015; Wang et al., 2018). The fact that, *in vitro,* RecA can catalyze an inverse strand exchange reaction with DNA or RNA thus promoting the assimilation of a transcript into duplex DNA (Kasahara et al., 2000; Zaitsev & Kowalczykowski, 2000) does not argue that this is the case for unscheduled recombinogenic R loops *in vivo*. More likely, the biological significance of this process relies on its use for replication initiation of prokaryotic cells as originally proposed (Zaitsev & Kowalczykowski, 2000), for replication-dependent recombination to restart stalled forks (Pomerantz & O’Donnell, 2008) or even for transcription-induced origin-independent replication (Stuckey, Garcia-Rodriguez, Aguilera, & Wellinger, 2015). Hence, DNA:RNA hybridization could occur *in trans* under regulated conditions but not spontaneously as unscheduled and harmful structures that would put genome integrity into risk. Thus, the assimilation of a transcript into a duplex DNA *in trans* would be tightly regulated and limited to specific reactions such as the case of telomerase or CRISPR and possibly other proteins yet to be discovered. For other cases, such as that of the GADP45 factor that binds to promoters harboring hybrids formed by lncRNAs (Arab et al., 2019), it is unclear whether such hybrids are formed *in trans* and in a GADP45-dependent manner.

Altogether, our results suggest that RNAs do not form hybrids *in trans*, so that the previously reported induction of Rad51-dependent ectopic genetic instability would be explained by R loop-mediated DNA breaks *in cis*.

## MATERIALS AND METHODS

### Yeast strains and Plasmids

Strains used were the wild-type W303-1A *(MATa ade2-1 can1-100 his3-11,15 leu2-3,112 trp1-1 ura3-1)* and its isogenic *hpr1*Δ::*HIS3* mutant U678-1C (*MAT*a) and U678-4C (*MATα*), *mft1*Δ::*KANMX* mutant (WMK.1A) (Chavez & Aguilera, 1997), *rnh1*Δ::*KANMX rnh201*Δ::*KANMX* (RNH-R), *rad51*Δ (WSR51.4A) (Gonzalez-Barrera, Garcia-Rubio, & Aguilera, 2002), *rad51*Δ::*HPH* (AYBY-7B), *hpr1*Δ*HIS3 rad51* Δ::*HPHMX* (AYBY-9B) and *hpr1*Δ *rad51*Δ (HPR51.15A) from this study. *rnh1*Δ::*KAN rnh201*Δ::*KAN* (HRN2.8A) and the wild-type HRN2.8A were from (Huertas & Aguilera, 2003). Wild-type (WGLZN) and *hpr1*Δ*HIS3* mutant with the GL-*LacZ::NATMX* inserted at the *LEU2* locus in Chromosome III were made in this study.

Yeast plasmids pCM179, pCM184, pCM189 and pCM190 were previously published (Gari, Piedrafita, Aldea, & Herrero, 1997). pRS314-GL-*LacZ* (Piruat & Aguilera, 1998) and pRS314-GL-*LacZi*, containing the recombination system with the *BamH*I fragment containing the *LacZ* sequence from pPZ (Straka & Horz, 1991), inserted in both sense and antisense orientations respect to the promoter, respectively, into the *Bgl*II site of pRS314GLB (Piruat & Aguilera, 1998), located between the two leu2 repeats. pCM184:AID was built by inserting the AID ORF from pCM189:AID (Santos-Pereira et al., 2013) into the *Not*I site of pCM190. pCM190-tet::LacZ400 was built by cloning the *Kpn*I-*Bam*HI 400-bp fragment of the 3’ from the *LacZ* gene into *Kpn*I-*Bam*HI digested pCM190. Plasmids pCM189:RNH1 (Castellano-Pozo, Garcia-Muse, & Aguilera, 2012), pCM189:AID, pCM184:RNH1 (Santos-Pereira et al., 2013), pRS316-LYΔNS (Prado et al., 1997) and pWJ1344 (Lisby et al., 2001) were also previously published.

### Yeast transformation

Yeast transformation was performed using the lithium acetate method as previously described (Gietz, Schiestl, Willems, & Woods, 1995).

### Recombination assays

Recombination frequencies were calculated as previously described (Gomez-Gonzalez, Ruiz, & Aguilera, 2011) as means of at least 3 median frequencies obtained each from 6 independent colonies isolated in the appropriate SC medium for the selection of the required plasmids. Recombinants were obtained by platting appropriate dilutions in selective medium. To calculate total number of cells, plates with the same requirements as for the original transformation were used. All plates were grown for 3-4 days at 30°C.

### Detection of Rad52-YFP foci

Spontaneous Rad52-YFP foci from mid-log growing cells carrying plasmid pWJ1344 were visualized and counted by fluorescence microscopy in a Leica DC 350F microscope, as previously described (Lisby et al., 2001). More than 200 S/G2 cells where inspected for each experimental replica.

### S9.6 immunofluorescence of yeast chromosome spreads

The procedure performed is similar to (Chan et al., 2014) with some modifications. Briefly, mid-log cultures (OD600= 0.5-0.8) were grown at 30°C; 10 ml of them were collected, washed in cold spheroplasting buffer (1.2 M sorbitol, 0.1 M potassium phosphate and 0.5 MgCl2 at pH 7) and then digested by adding at the same buffer 10 mM DTT and 150 mg/ml of Zymoliase 20T. The digestion was performed during 10 minutes (37°C) and stopped by mixing the samples with the solution 2 (0.1 M MES, 1M sorbitol, 1mM EDTA, 0.5 mM MgCl_2_, pH 6.4. Later, spheroplasts were centrifuged carefully 8 minutes at 800 rpm, lysed with 1% vol/vol Lipsol and fixed on slides using Fixative solution (4% paraformaldehide/3.4% sucrose). The spreading was carried out using a glass rod and the slides were dried from 2 hours to overnight in the extraction hood.

For the immuno-staining, the slides were first washed in PBS 1X in coplin jars and then blocked in blocking buffer (5% BSA, 0.2% milk in PBS 1X) over 10 minutes in humid chambers. Afterwards, slides were incubated with the primary monoclonal antibody S9.6 (1 mg/ml) in a humid chamber 1 hour at 23°C. After washing the slides with PBS 1X for 10 minutes, the slides were incubated 1 hour at 23°C in the dark with the secondary antibody Cy3 conjugated goat antimouse (Jackson laboratories, #115-165-003) diluted 1:1000 in blocking buffer. Finally, the slides were mounted with 50 μl of Vectashield (Vector laboratories, CA) with 1X DAPI and sealed with nail polish. More than 300 nuclei were visualized and counted to obtain the fraction of nuclei with DNA:RNA hybrids.

## ACKNOWLEDGMENTS

Research was supported by the Spanish Ministry of Economy and Competitiveness (BFU2016-75058-P) and the European Union (FEDER). B.G.-G. was funded by a grant from the Spanish Association Against Cancer (AECC).

## COMPETING INTERESTS

The authors declare no competing interests.

## Notes

### Competing Interest Statement

The authors have declared no competing interest.

